# Construction of a neural network energy function for protein physics

**DOI:** 10.1101/2021.04.26.441401

**Authors:** Huan Yang, Zhaoping Xiong, Francesco Zonta

**Author notes:** For correspondence, please contact Francesco Zonta or Huan Yang.

## Abstract

Classical potentials are widely used to describe protein physics, due to their simplicity and accuracy, but they are continuously challenged as real applications become more demanding with time. Deep neural networks could help generating alternative ways of describing protein physics. Here we propose an unsupervised learning method to derive a neural network energy function for proteins. The energy function is a probability density model learned from plenty of 3D local structures which have been extensively explored by evolution. We tested this model on a few applications (assessment of protein structures, protein dynamics and protein sequence design), showing that the neural network can correctly recognize patterns in protein structures. In other words, the neural network learned some aspects of protein physics from experimental data.

## Introduction

The best possible description of protein molecules should involve Quantum Mechanics (QM) and the solution of the Schrödinger equation. However, in the study of biological systems, models based on approximate classical potential have been successful to understand how proteins and other biological molecules interact. Two main categories of simplified potentials have emerged to describe proteins: physics-based potentials and knowledge-based potentials. Both methods rely on physical intuition to map interactions between atoms into simple functional forms that depend on various parameters. In the first case, the parameters are derived by comparison with high precision QM calculations or experiments. These potentials take often the name of force fields and are mainly used to perform molecular dynamics (MD) simulations of biological systems (*1*–*3*). In the second case, probability distribution of observables (e.g. distances, angles, native contacts etc.) are directly obtained from experimental information and transformed into a statistical potential (*4, 5*). These potentials are mainly used for applications that are too much computationally expensive for MD simulations. The two approaches are not antithetic and their boundary is blurred as many hybrid potentials are developed (*6, 7*).

Approximate classical potential, however, still present some limitations that goes in two opposite directions. For some applications, their simple functional form is not able to consider all the details required for an accurate description of the system, meanwhile, in other cases they are too computationally complex and cannot reach time and size scales directly comparable with experiments. Although QM calculations and hybrid methods (*8*–*10*) can be used to improve the accuracy of the calculations, and enhanced sampling methods (*11*) or coarse grained force fields (*12, 13*) can be used to scale up in both time and size scales, the core problems still exist.

Deep neural networks (*14*) are emerging as new ways to approximate complex physical energy functions and opening new opportunities to compute molecular properties accurately and fast. In this approach, the energy function has not a fixed functional form, but is represented by a neural network. This strategy has been applied to fit the potential energy surface calculated with density functional theory for small molecules (*15, 16*). More recent works have developed neural network energy functions for multi-scale modeling of molecules following a bottom-up approach, e.g. fitting potentials at atomic resolution to QM calculations (*17*), or fit coarse grained potentials to atomic molecular dynamics simulations (*18*). These energy functions are still not transferable to general proteins but could lead to a paradigm shift in the field of force fields in the future.

In this paper we use an unsupervised deep learning method to construct a neural network energy function for proteins, using a top-down approach, i.e., from experimental data or protein structures and sequences. We want to construct an energy function that is general enough to be applied to various tasks in pair with other standard methodology (for example to run MD simulations, or discriminate decoys from native configurations). The energy function should depend only on the protein configuration, be time independent and differentiable, possess rotational and translational invariance, so that it could correctly reproduce protein physics.

### Design of the neural network energy function based on protein building blocks

In the construction of a data-driven energy function for proteins, the idea is to use data sampled by evolution to approximate the physical energy landscape, because evolution has sampled a repertoire of diverse proteins by tinkering and reusing building blocks with low energies (*19*). The choice of the evolutionary data sets is critical and a naive training of the network with all the known experimental protein structures and sequences will likely fail to reproduce correctly protein physics. In an ideal case, we would like to have a large sample of unbiased sequence-structure pairs, but in reality, when looked from the point of view of their physical properties, the set of proteins with known experimental structures is both small and redundant. The known folding domains (*20, 21*) are incomplete i.e. structures that appear legitimate from an energetic point of view may have not yet been selected or explored by evolution (*22*). However, we can hypothesize that evolution had enough time to extensively explore the configuration space of smaller 3D local structures (*23*), and that this information is already contained in the available protein databases.

For this reason, we describe a protein as a collection of small-scale structures (building blocks) paired with their sequence information. In the training phase, the building blocks are selected from a non-redundant set of protein structures and their homologous sequences (see Fig. 1 and Methods). We assume that the set is complete, i.e. small-scale structures that are not represented in this set are improbable and have high energies. We then assume that the energy function of the whole protein is the sum of the contributions of each building block. The energy function obtained in this way will be local and addictive and, if trained properly, the neural network will be able to recognize alternative local minima, which corresponds to alternative configurations or active states etc.

**Fig. 1.**
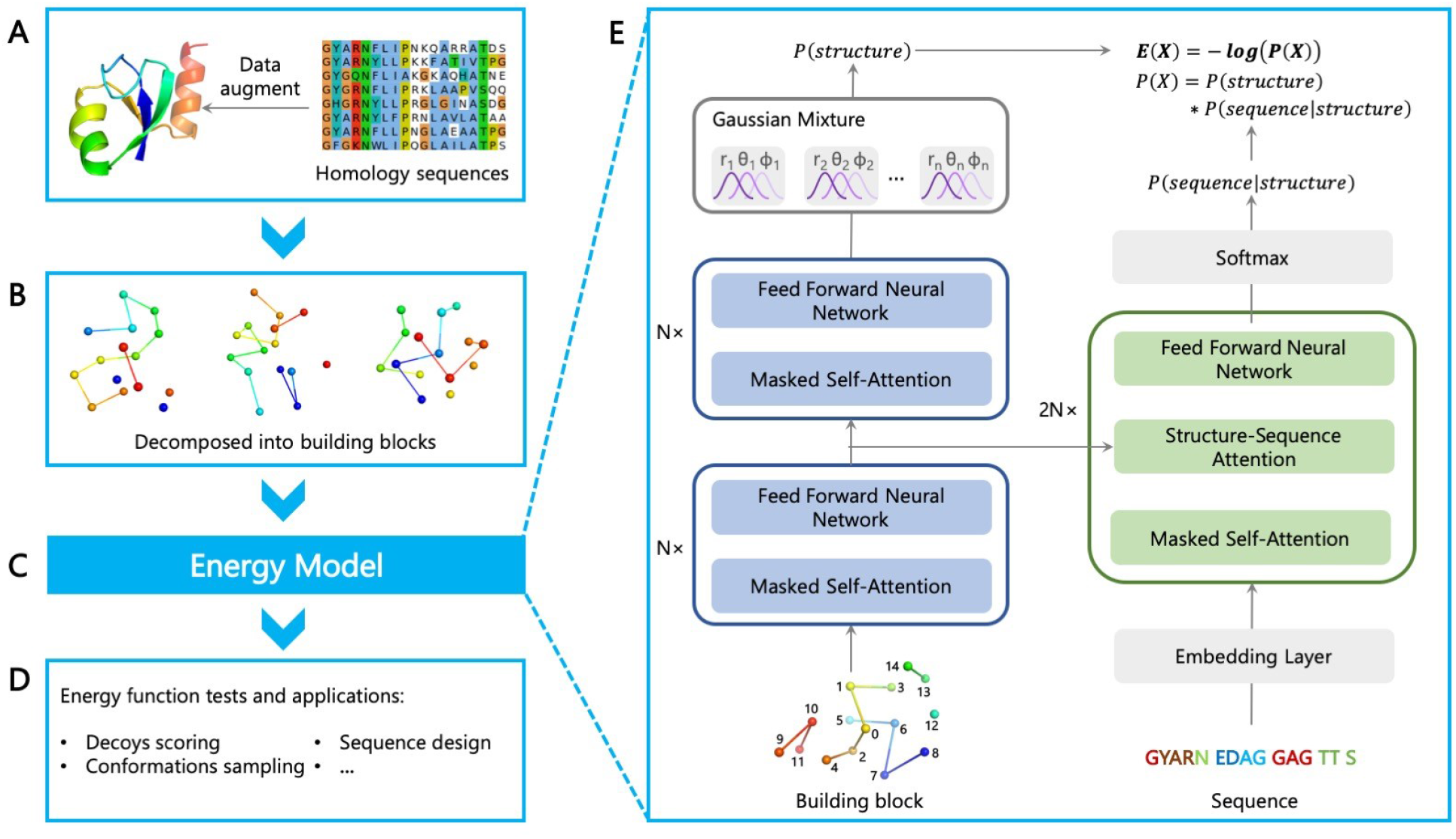
Overview of the NNEF model. (A) We use a sample of non-redundant protein structures and augment the training data with their homologous sequences. (B) The local structure around each residue defines the building block and is used as the input of the energy model. Each building block *X* includes the residue itself, its 4 nearest residues along the sequence and the 10 other nearest residues in the 3D space. Each residue is represented as one bead. (C) The energy model is illustrated in (E), where we fit a probability density function and calculate the energy as *E*(*X*) = −*logP*(*X*). The total energy of the protein is the sum of energies of all building blocks. In the auto-regressive model, we separate the structural and sequence features, and calculate *P*(*X*) = *p*(*Structure*(*X*)) · *p*(*Sequence*(*X*) | *Structure*(*X*)) using the transformer network architecture. We use *softmax* function to calculate the probabilities of discrete features and use Gaussian mixtures to calculate the probabilities of continuous features. (D) The learned energy function can be applied for various tasks, such as decoys scoring, conformations sampling, and sequence design without explicitly trained for any of those.

#### Protein building blocks

Definition and representation of building blocks rely on human intuitions. They can be pairs of residues, peptide fragments, groups of residues, etc. They can be represented by atoms, virtual beads, and various geometric and/or chemical features. Furthermore, the size of the building blocks should be small enough in order to minimize the risk of learning from under sampled sets, but large enough, so that they can represent complex tertiary structures.

In our current model, we define a building block as the local structure of 15 amino acids around a residue. This includes the residue itself, its 4 nearest residues along the sequence and the 10 other nearest residues in the 3D space (Fig. 1B and fig. S2). With this definition, building blocks are typically composed by a few noncontiguous segments. As we will show, the energy function can be used to run MD simulations, and in this particular case the building blocks change dynamically with time. To represent the structure of building blocks, we use very simple low resolution geometric features i.e., the coordinates of beads at the *C*_*β*_ positions (*C*_*α*_ for Glycine), and the connectivity of these beads based on the protein sequence. More geometric features, such as the positions of backbone atoms, the side chain positions, and hydrogen bonds can be added as needed. To make the energies rotational-translational invariant (as it should be for a Hamiltonian function), the coordinates of all the beads composing each building block are rotated to the same local internal coordinate system (fig. S1).

#### Machine learning model

After decomposing the proteins into building blocks and extracting their features, we characterize the evolutionary landscape of building blocks by fitting a probability density function, and set for each building block *X, E*(*X*) = −*logP*(*X*). The total energy of a protein *Y* is the sum of the energies of all building blocks composing 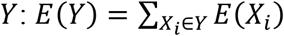. The probability function *P*(*X*) is obtained with an auto-regressive model and maximum-likelihood training from a set of *N* building blocks {*X*}. Each building block *X* can be viewed as a list of k variables *x*_1_, *x*_2_, …, *x*_*k*_ ordered in a given scheme, and its probability *P*(*X*) can be expanded according to the Bayesian rule as *P*(*X*) = *Π*_*j*_*p*(*x*_*j*_|*x*_<*j*_). In the ordering scheme, we separate the structural and sequence features, so that *P*(*X*) = *p*(*Structure*(*X*)) · *p*(*Sequence*(*X*) | *Structure*(*X*)). With *P*(*X*) expanded as a chain of conditional probability functions, each conditional probability *p*(*x*_*j*_|*x*_<*j*_) is represented as a neural network that shares parameters with other conditional probability functions. The neural network architecture of the auto-regressive model is shown in Fig. 1E and detailed in the Methods section.

### Understanding the neural network energy function

To understand whether the Neural Network Energy Function (NNEF) is able to grasp correctly the protein physics, we test it in several tasks described hereafter.

#### Scoring decoys generated by modifications of the native structures

A good energy function should be able to discriminate native-like from non-native conformations. We tested the energy function against the 3DRobot decoy set (*24*) and a second set of decoys that we generated by sampling normal modes of protein structures. The 3DRobot decoy set includes 200 non-homologous single domain proteins (48 in the *alpha* class, 40 in the *beta* class, and 112 in the *alpha/beta* class) and 300 structural decoys for each of these proteins, with root-mean-square deviation (RMSD) ranging from 0 to 12 Å. The second set was generated for a test sample of 18 small (<120 residues) proteins which comprises 4 *alpha* proteins, 7 *beta* proteins, and 7 *alpha/beta* proteins. In all proteins of both decoy sets, the energy tends to increase with the distance from the native, with native-like decoys having low energies and decoys with large RMSD having high energies (a typical result for each set is shown in Fig. 2).

**Fig. 2.**
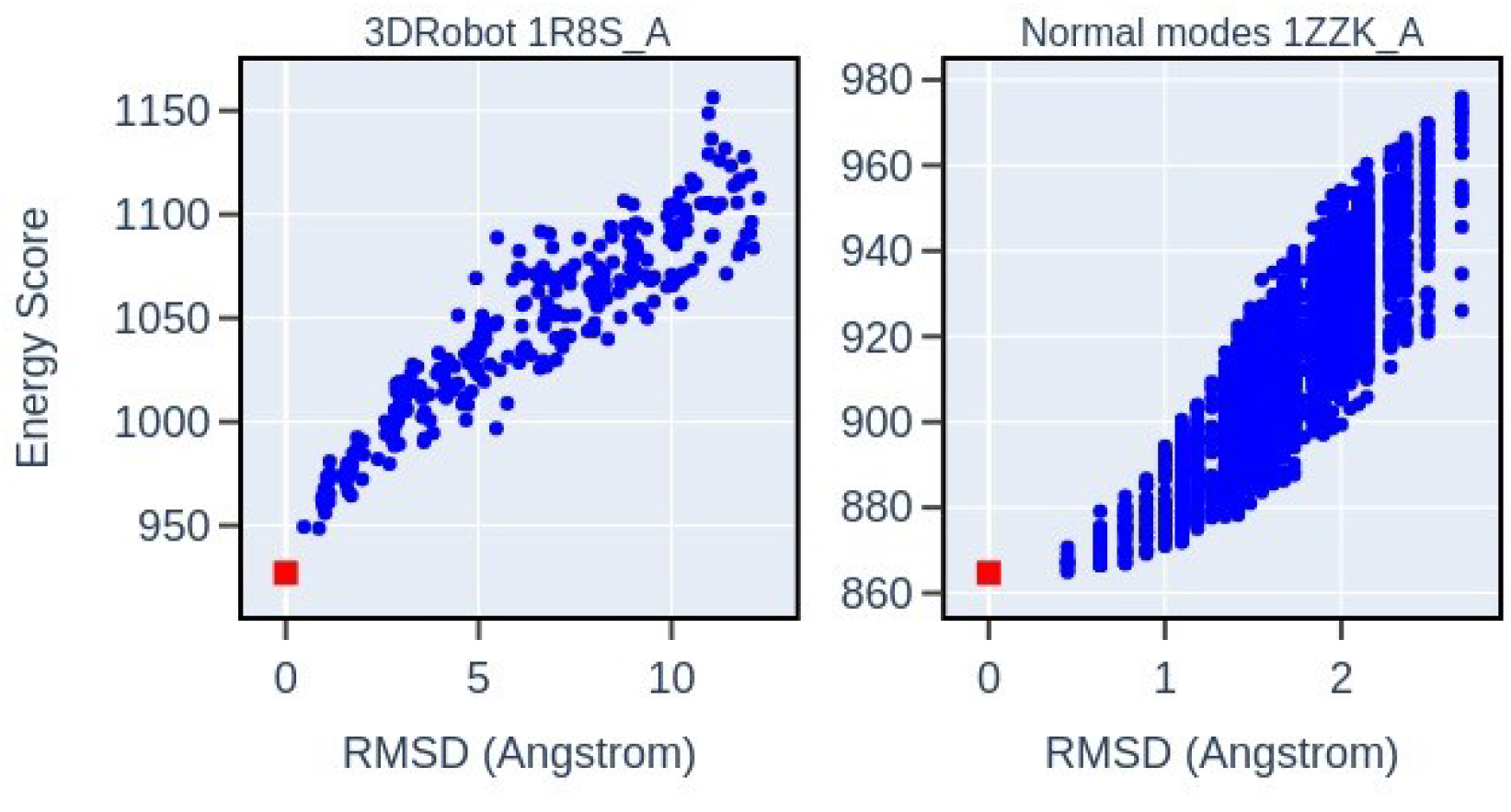
Scoring decoys generated by modifications of the native structures. The left panel shows one typical example protein in the 3DRobot decoy set. The right panel shows one typical example protein in the normal modes decoy set. Red square is the native structure and blue dots are decoys. In all proteins of both decoy sets, the energy increases with the distance from the native.

#### Scoring structure predictions

Another way in which we tested the quality of the NNEF is to score predictions generated in the 14th Critical Assessment of Techniques for Protein Structure Prediction (CASP14) contest. This is a bigger challenge than scoring the decoy sets generated by modifications of native structures, because such predicted structures cover more diverse conformations and are optimized according to some other scoring functions. Each of the 97 protein domains in CASP14 has about 200-500 structural models. We evaluate the NNEF for each of these predictions and measured its correlation with the CASP14 Global Distance Test Total Score (*GDT*_*TS*_), which measures the overall quality of the prediction. For about 70% of the sets, we obtain a Pearson correlation coefficients |*ρ*| > 0.75 (Fig. 3A, B). In the remaining cases (many of which are proteins that belong to complexes and for which their tertiary native structures could depend on the environment), the energy of the best CASP prediction is always near the global minimum of the evaluated NNEF for the whole set. In other words, the energy function appears to be quite successful in detecting good configurations but could be fooled by reasonable bad configurations (Fig. 3C, D).

**Fig. 3.**
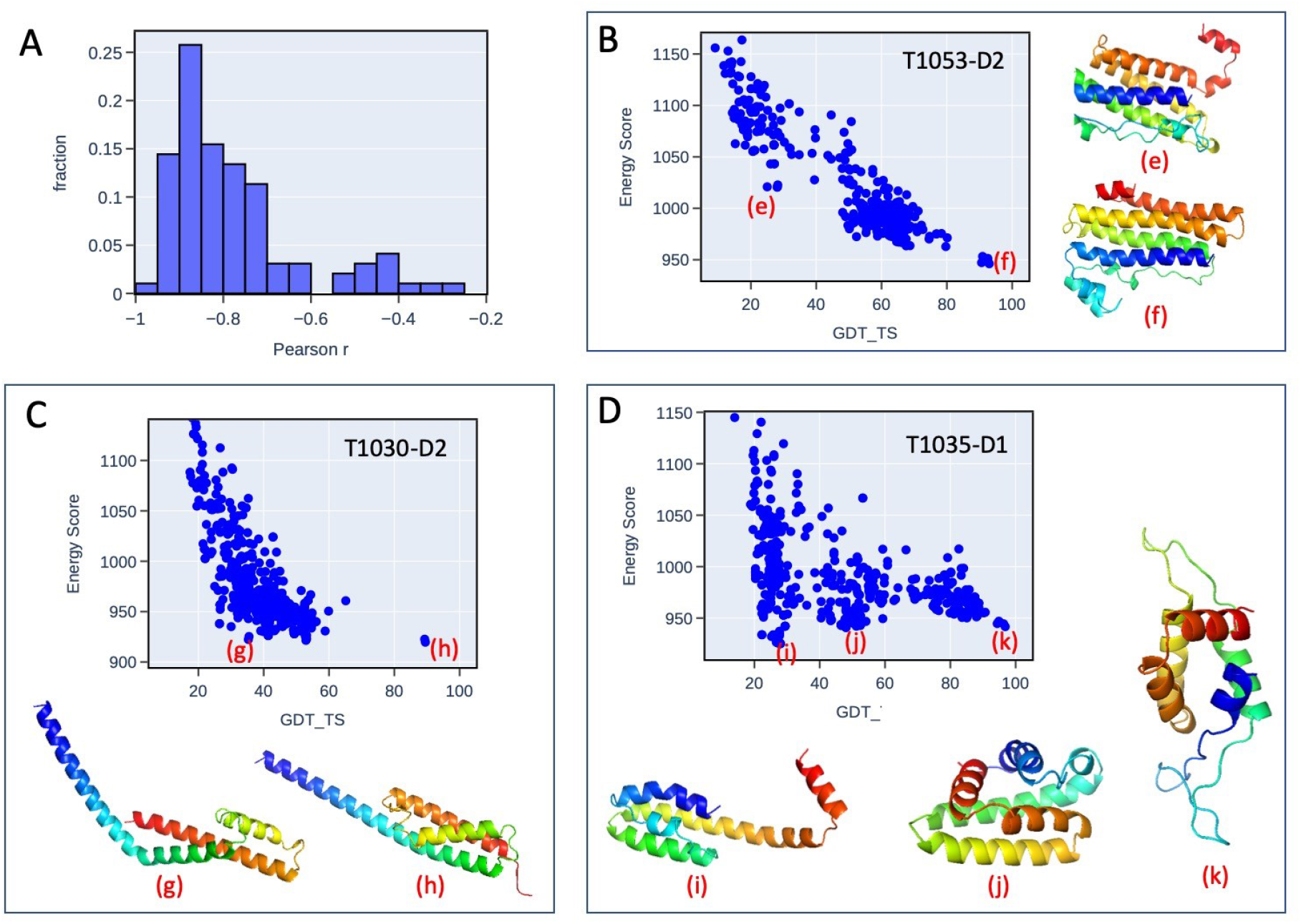
Scoring structure predictions in CASP14. Panel (A) shows the histogram of Pearson correlation coefficients *ρ* between the energy score and CASP GDT_TS score for proteins in CASP14. About 70% of proteins have Pearson correlation coefficients |*ρ*| > 0.75. Panels B, C, and D show three particular cases. 3D structures of some decoys are shown with indicators of their positions in the plot of energy vs. GDT_TS. Panel (B) shows an example of good correlation between the NNEF energy and CASP GDT_TS score.. Panel (C) shows an example case of a protein with simple *alpha* helices fold. In this case, some models with non-native helix organizations have comparable energies to native-like models. Panel (D) shows an example case of a protein involved in complex context. Some models with wrong folds have energies lower than the native-like models.

#### Evaluating the energy of configurations within a MD trajectory

The scoring of decoys and model predictions indicate that the learned energy function can be generalized to non-native configurations, despite having been trained only with native structures. To further explore this feature, we use the NNEF to score a conformation ensemble of a small protein (Fip35) sampled over a 100-microsecond long MD trajectory (*3*). In this time window Fip35 explores regions of the phase space far from the native configuration and undergoes multiple folding and unfolding events. We compare the RMSD and the energy evaluated with NNEF along the MD trajectory. As shown in Fig. 4, the folded states have low energy scores, while the unfold states generally have higher energy scores, which further shows the energy function can generalize to non-native conformations.

**Fig. 4.**
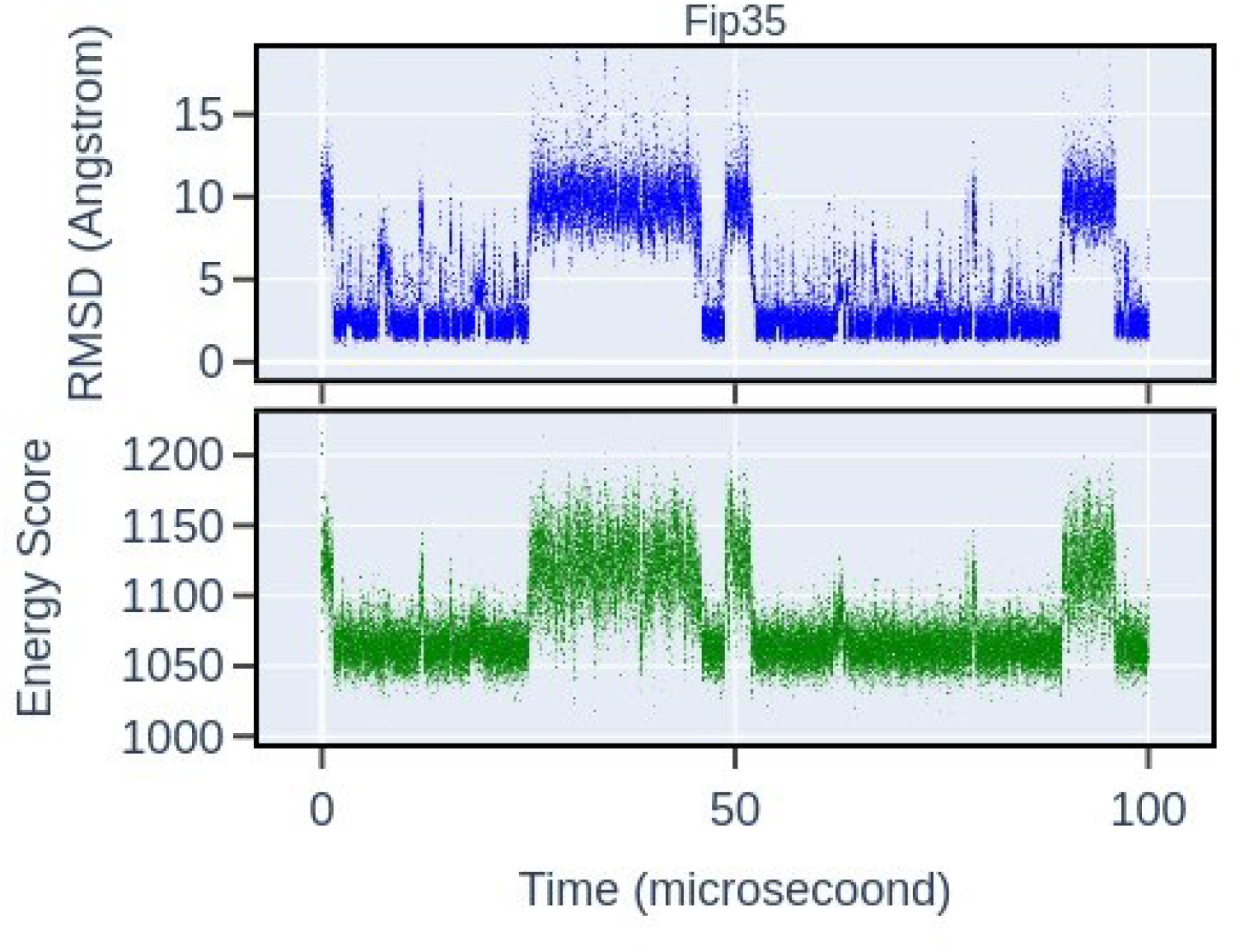
Evaluating the energy of configurations within a MD trajectory for a small protein Fip35. The protein undergoes multiple folding and unfolding events. The RMSD and the energy along the MD trajectory are in a good correlation, suggesting the energy function can generalize to non-native conformations.

#### Performing MD simulations

The energy function can be interpreted as a Hamiltonian function, and can be used to study protein dynamics with an implicit solvent (Langevin dynamics). In the dynamics, the Cartesian coordinates 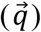 of the residue beads are updated at each step as 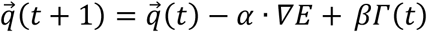. *∇E* +*βΓ*(*t*), where *E* is the NNEF for the protein at time *t* and *Γ*(*t*) is a Gaussian noise with null average and unitary variance. The coefficients *α* and *β* are related to the physical friction coefficients, the integration time steps, the temperature of the system, and the physical units of energy values. With a proper choice of their values (see Methods) we can observe that the dynamics generated by this method produce protein fluctuations consistent with those found with classical MD based on force fields. After fixing values for *α* and *β*, we generated a 30000 steps Langevin dynamics for the test set of 18 small proteins and compared it to MD simulations obtained with *amber14SB* force field (*2*) using *openMM* (*25*). We can observe that the root-mean-square fluctuations (RMSFs) calculated for each residue across the whole set of protein obtained by the two methods have comparable distribution, and their difference stays lower than 0.5 Å in most cases. For most proteins in the sample, the two dynamics produce RMSFs in good correlation across the whole sequence (see Fig. 5 and fig. S5). A complete assessment of the validity of the NNEF as a force field deserves a much more exhaustive analysis, which is out of the scope of this article, however, these preliminary results indicate that the NNEF can understand how a protein should behave when it is excited by thermal fluctuation.

**Fig. 5.**
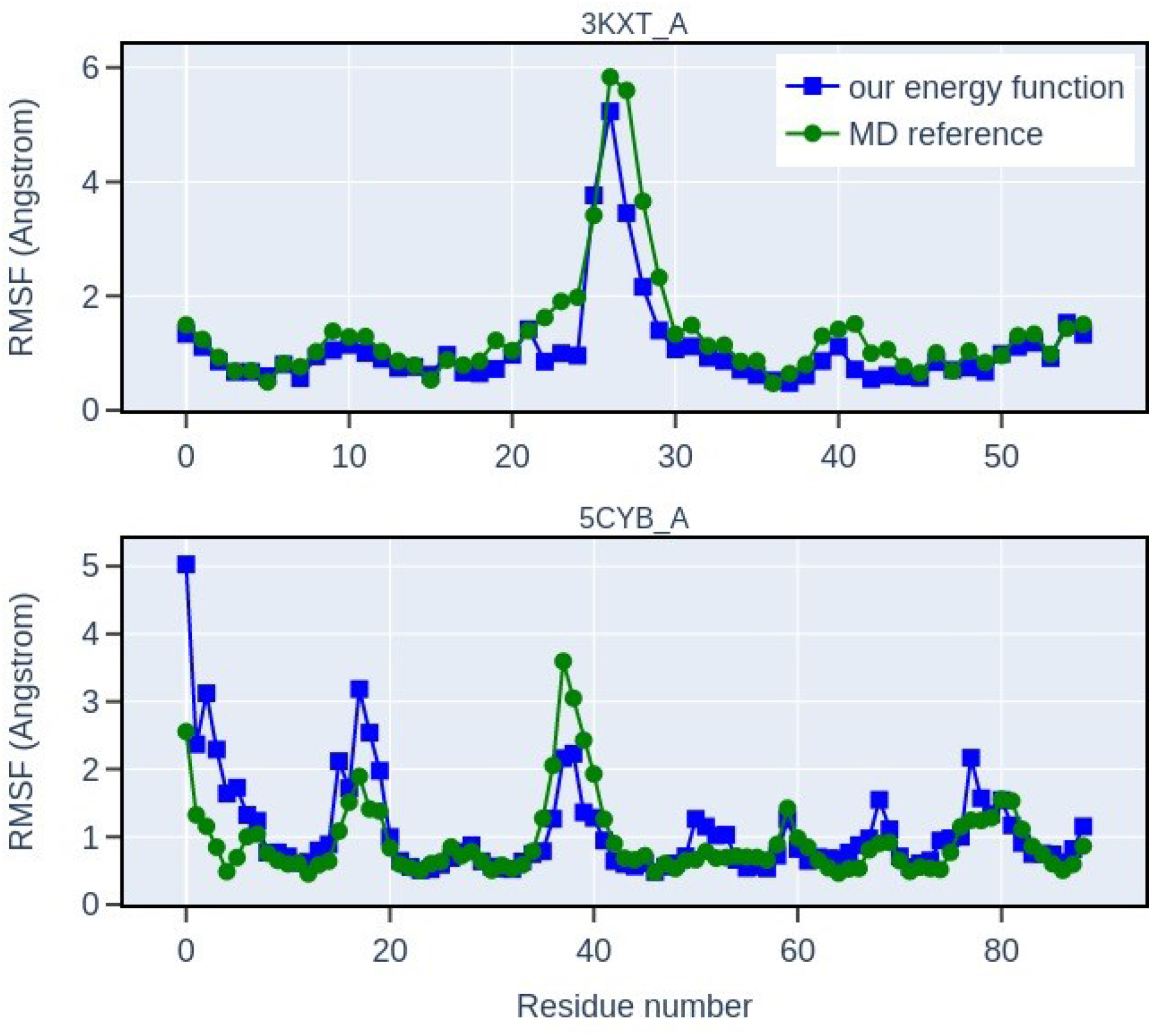
RMSFs resulted from Langevin dynamics simulations using the NNEF. We compare the RMSFs of the trajectories from the NNEF (blue squares) with the RMSFs of the trajectories from classical MD simulations obtained with *amber14SB* force field (green dots). Two examples with high correlation are shown here. For most proteins in the sample, the two dynamics produce RMSFs in good correlation across the whole sequence (see also fig. S5).

#### Importance of the sequence in the energy function

The next question we want to address is whether the energy function have learned the difference between the various amino acids or if it only learned local structural properties of a protein chain. This is not a given, since it is possible to construct protein models that correctly describe local pattern without any sequence information (*26*). To investigate this point, we fix the structure and evaluate the energy of a sample of 100 proteins after making changes to its sequence. The “decoy” sequences were generated in four different ways: (1) substituting residues with chemically similar residues, (2) shuffling the residues in the sequence, (3) mutating residues to random ones, (4) mutating all residues to the same residue type. In most of the cases, the energies of the mutated sequences are higher than the native ones, and random mutations produce higher energies than mutations to similar amino acids (fig. S3). However, sequences that are rich in alanine appear to be favored by the network and almost all poly-alanine decoys have a better energy than the real sequence (fig. S4). This happens, to a lesser extent, also to poly-leucine and poly-histidine, but not for the other amino acid.

#### Protein sequence design

To further explore the interplay between sequence and structure in the NNEF, we redesigned the full sequences from their backbone structures for the test sample of 18 small proteins. Starting from a random sequence, we run simulated annealing to obtain a sequence that should fold to the desired structure. At each step of the annealing process, we propose a random point mutation for the protein sequence. The mutation is accepted or rejected according to a Metropolis algorithm. In most of the cases, the simulations converge to sequences having energies lower than the native after a few thousands of steps. We designed in this way 100 sequences for each target protein within the mentioned test sample. Amino acid frequencies for these 1800 designed sequences are shown in Fig. 6A. As we can observe, the annealing process converges to sequences that favor 10 amino acids (Ala, Val, Leu, Gly, Pro, Ser, Thr, Arg, Glu, and Asp). These 10 amino acids have relatively high frequencies in natural protein sequences, and may be the first 10 that joined biological proteins early in evolution according to the theory on temporal order of amino acids in evolution (*27*). Overall, the average sequence recovery fraction is about 25% for the total sample of 1800 designed sequences. We also analyzed the differences between core and surface residues of the designed sequences (Fig. 6B). Core residues are mostly hydrophobic (Ala, Val, Leu), while the polar residues are more likely to be exposed. Moreover, core residues are more conserved compared to surface residues, when considering all the 100 designed sequences for each structure. Finally, to give a measure of the reliability of the protein design method, we selected two random designed sequences for each protein and predicted their structures using *TrRosetta* (*28*). For about two thirds of the designed sequences, the predicted structures match the target structures well (Fig. 6C and fig. S6).

**Fig. 6.**
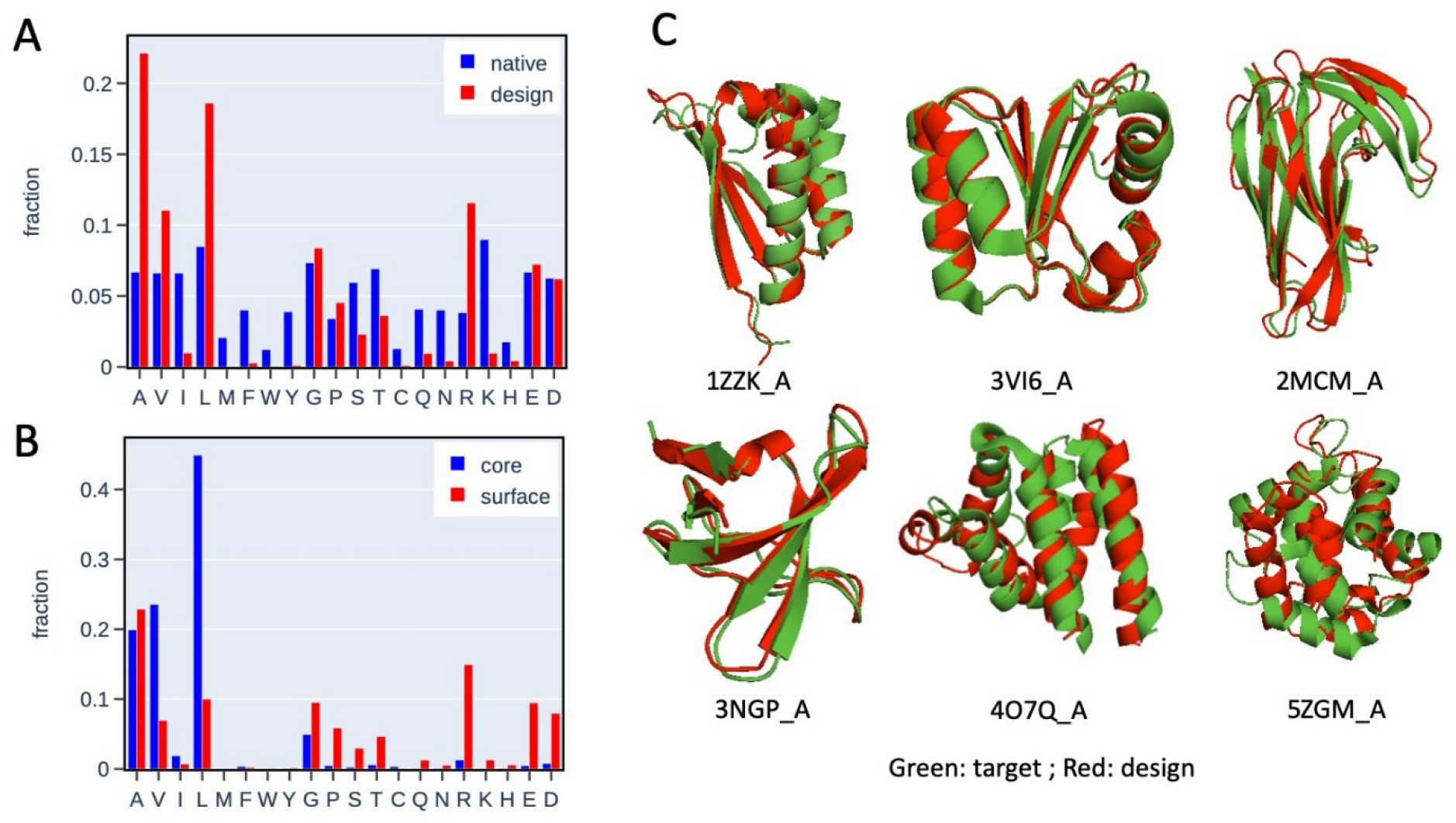
Results of protein full sequence design given the backbone structure. (A) Amino acids frequencies of the 1800 designed sequences and the native sequences for the test sample of 18 proteins (100 sequences for each protein). Designed sequences show preference for 10 amino acids (Ala, Val, Leu, Gly, Pro, Ser, Thr, Arg, Glu, and Asp). (B) Amino acids frequencies of the residues in the core and surface of the designed proteins. Core residues are mostly hydrophobic (Ala, Val, Leu), while the polar residues are mostly in the surface. (C) A few examples of the target structures and predicted structures using *TrRosetta*. For about two thirds of the designed sequences, the predicted structures can match the target structures.

## Discussion

In this paper we explored the idea of using a neural network energy function to describe Protein physics without explicitly fixing its functional form. We describe the local environment of each residue as a building block that takes into account also residues far in the sequence but near in the 3D structure, and hypothesize that evolution had enough time to extensively explore the configuration space of building blocks. Then we use unsupervised machine learning methods to learn a NNEF from the structure and sequence data of building blocks. Our results indicate that the learned NNEF is general and transferrable to various problems (albeit with various degree of success), without being aimed at any of those. This suggests that the network has learned some physical characteristics of local patterns in proteins.

### Comparison to related works

Deep learning has been widely used to model proteins in the past. However, the approach presented here is taking a different point of view compared with previous studies. In many studies, a scientific question is translated into a specific supervised learning task. While from the unified perspective of energy based model (*29*), all these supervised models can be viewed as energy functions, they are usually only applicable to the particular task for which they are trained and they are not required to be a realistic approximation of the physical energy function of proteins.

Recent works have shown that it is possible to successfully predict protein structures using deep learning methods (*28, 30*). These models could have implicitly learned some aspects of protein physics. For example, deep learning models trained to predict protein structures from multiple sequence alignments can also be used to design protein sequences and evaluate protein stability (*31, 32*). The generality and transferability of these models still need further studies.

A different approach has been to exploit the massive number of sequences, and apply unsupervised methods, developed for natural language processing, to protein sequences (*33*–*35*). The learned energy functions can be used to predict secondary structure and contacts in tertiary structure. The discrete sequence data is very sparse and makes this approach very difficult. Adding structures to the learning can make the model more data efficient.

### Moving forward from here

Neural network energy function (or energy-based model) is very flexible, and our current implementation is far from realizing its full potential. We can improve the method by extensively testing different ways to combine structures and sequences, different representations of the proteins and building blocks, and better training methods and loss functions. It is also possible to refine the parameters of the energy function by supervised fitting to multiple tasks, after the unsupervised training. Furthermore, the always increasing availability of experimental data could also lead to better NNEF in the future.

Another important advantage, compared to more traditional methods to parameterize an energy function, is that we are not restrained to a single resolution. The protein description could be fully atomistic, coarse grained, or mixed, without any loss of generality. The network can learn how different resolutions should merge. It is foreseeable to use a neural network energy function to build simulations in which some parts have the desired level of details and other less interesting parts are at a lower resolution.

## Conclusions

Is it possible for a neural network to understand protein physics from biological examples? In this paper we used unsupervised deep learning methods to derive a neural network energy function for describing amino acids interactions within proteins. The learned energy function can discriminate decoys, assess qualities of structural models, sample structural conformations, and design new protein sequences. Altogether, this suggests that a high-level approximation of protein physics can be learned from data and this methodology can lead to a completely new way to parameterize protein energy function.

## Materials and methods

### Structure and sequence dataset

As explained in the results section, to train the network we want to curate a non-redundant sample of structures and associate each structure to a family of homologous sequences. To reduce the redundancy in the structures, we use a sample of *PDB* chains in a version of the *PISCES CulledPDB* (*36*), in which the percentage identity cutoff is 50%, the resolution cutoff is 3.0 Å, the R-factor cutoff is 1.0, and the total number of chains is about 29,000. Only structures solved by X-ray crystallography are used. Then, all the chains are matched to the *HH-suite PDB70* database (*37*) to get the aligned sequence data. The number of matched chains is about 19,000. We filter the aligned sequences using *hhfilter* by requiring <50% sequence identity to the *PDB* sequence and >50% sequence coverage. After the filtering, we generate homologous structures by simply mapping the aligned sequences to the coordinates of the structure. Given a sequence A in a *PDB* chain and an aligned homologous sequence B, we ignore insertions in sequence B, and substitute gaps in sequence B with the aligned sites in sequence A. In this way, the resulting chimeric sequence has the same length as sequence A and can be mapped to the coordinates of A.

To test the transferability of the energy function, we use a radical partition of the dataset. All chains are matched to the structural classifications in the CATH 4.2 database (*20*). Each chain can include more than one CATH domains. In the classification, a chain is classified as one class (e.g., *alpha/beta*) if all the CATH domains in the chain are classified as that class (*alpha/beta*). The training data includes only the *alpha/beta* chains. We get a total of about 7500 *alpha/beta* chains, and use 7000 chains as the training dataset, and 500 chains as the validation dataset. The trained energy function is tested on all protein classes, including *alpha/beta*, mainly *alpha*, and mainly *beta* proteins.

### Neural network model

#### Building blocks

To make the energies rotational-translational invariant the coordinates are rotated to the same local internal coordinate system of the central residue (fig. S1). The coordinate system is defined so that the residue *a* − 1is on X-axis, and the residues *a* − 1and *a* + 1are on X-Y plane. The Cartesian coordinates of all beads are then converted to Polar coordinates.

#### Ordering scheme in the autoregressive model

In the autoregressive model, the variables in both the structural and sequence features are ordered based on the same beads order (fig. S2), according to the following rule. Each building block X can be viewed as a graph with the central bead *a* as the root node. The central segment is visited first with the order (*a, a* − 1, *a* + 1, *a* − 2, *a* + 2). Then the other segments are visited in an order of increasing distance to the central bead. Within each surrounding segment, the beads are visited in an order of increasing primary sequence number.

#### Neural network architecture

The neural network architecture of the auto-regressive model is shown in Fig. 1E. The input data includes the coordinates, the bond connections, and the residue types of the 15 residues belonging to a building block. Bond connections, residue types, and positional labels 1−15 are then converted to vectors by learned embeddings. After this step the coordinates, bond connection vectors and positional vectors are concatenated as structural features, while residue types and positional labels are concatenated as sequence features. The encoder for structural features has 4 standard transformer encoder layers. The outputs after 2 encoder layers are used as the latent codes of the structural features and passed to the decoder. The decoder for sequence features has 4 standard transformer decoder layers. Both the encoder and decoder layers use the standard transformer layer (*38*). The attentions are masked by position-based causal masks, i.e., each position can only pay attention to positions before it.

#### Neural network training

For the predictions of the discrete labels, such as the bond connections and the residue types, we use *softmax* function to get the probabilities. For the predictions of the continuous variables, such as the radius and angles in the coordinates, we calculate the probabilities using mixtures of Gaussian functions: *p*(*x*) = ∑_*j*_*c*_*j*_*G*(*x, μ*_*j*_, *σ*_*j*_), where *G*(*x, μ*_*j*_, *σ*_*j*_) is a Gaussian function of the variable *x* with average *μ*_*j*_ and variance *σ*_*j*_, and *c*_*j*_, *μ*_*j*_, *σ*_*j*_ are outputs of the neural network. Thus, the network calculates the conditional probabilities of the next residue’s coordinate given the coordinates of previous residues. It also calculates the conditional probabilities of the next residue type given previous residue types and the coordinates of all residues. After getting the conditional probabilities, we calculate the energy terms as the negative *log* of the probabilities. The energy of the building block is the weighted sum of all the energy terms. We train the network to minimize the energies, i.e., the loss function is simply the energy values. During training, we use mini-batch training, Adam optimizer with starting learning rate 5 ×10^−5^ and *betas* = (0.9, 0.99), and L2 regularization with weight 10^−6^.

### Langevin dynamics

In the results section, we showed that we can use the NNEF to run Langevin dynamics, according to the following equation: 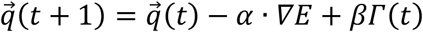. *∇E* +*βΓ*(*t*), where *E* is the NNEF for the protein at time *t* and *Γ*(*t*) is a Gaussian noise with null average and unitary variance, *α* and *β* are physical parameters of the simulation. To decide proper values for these two parameters, we run a grid search using short simulations (*α* = [1e-3, 3e-3, 5e-3, 7e-3, 0.01, 0.015, 0.02, 0.04], *β* = [0.01, 0.03, 0.06, 0.1, 0.15]). When *α* and *β* are small, the dynamics is very slow and the residues in the protein are almost locked in the initial position. When *β* is large, the protein unfolds after a small number of time steps quickly. We decide to use fixed values in the middle (*α* = 0.01 and *β*=0.05) to run simulations.

*The code will be publicly available at http://github.com/lahplover/nnef/.*

## Supplementary Materials

PDB IDs of the 18 small proteins in the test sample: 1ZZK_A, 2MCM_A, 2VIM_A, 3FBL_A, 3IPZ_A, 3KXT_A, 3NGP_A, 3P0C_A, 3SNY_A, 3SOL_A, 3VI6_A, 4M1X_A, 4O7Q_A, 4QRL_A, 5CYB_A, 5JOE_A, 5ZGM_A, 6H8O_A

**Fig. S1.**
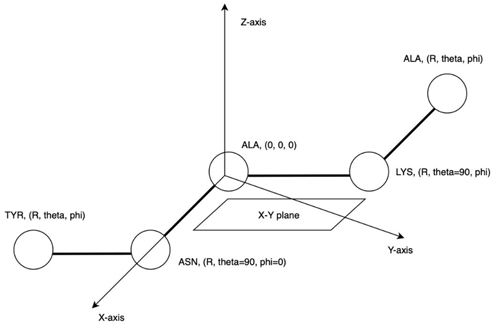
This figure shows the local internal coordinate system of a building block around a central residue. The coordinate system is defined so that the residue *a* − 1is on X-axis, and the residues *a* − 1and *a* + 1are on X-Y plane.

**Fig. S2.**
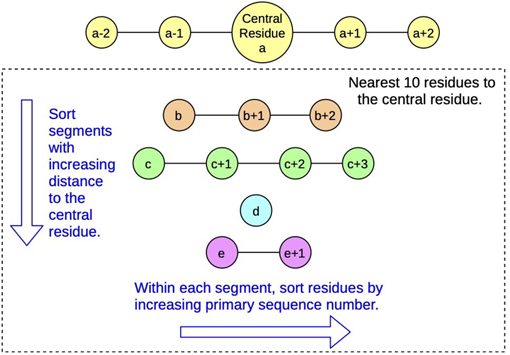
A typical building block has a central segment of 5 residues and a few other segments around the central residue. In the ordering scheme of the autoregressive model, the central segment is visited first with the order (*a, a*−1, *a* + 1, *a* − 2, *a* + 2). Then the other segments are visited in an order of increasing distance to the central residue. Within each surrounding segment, the residues are visited in an order of increasing primary sequence number.

**Fig. S3.**
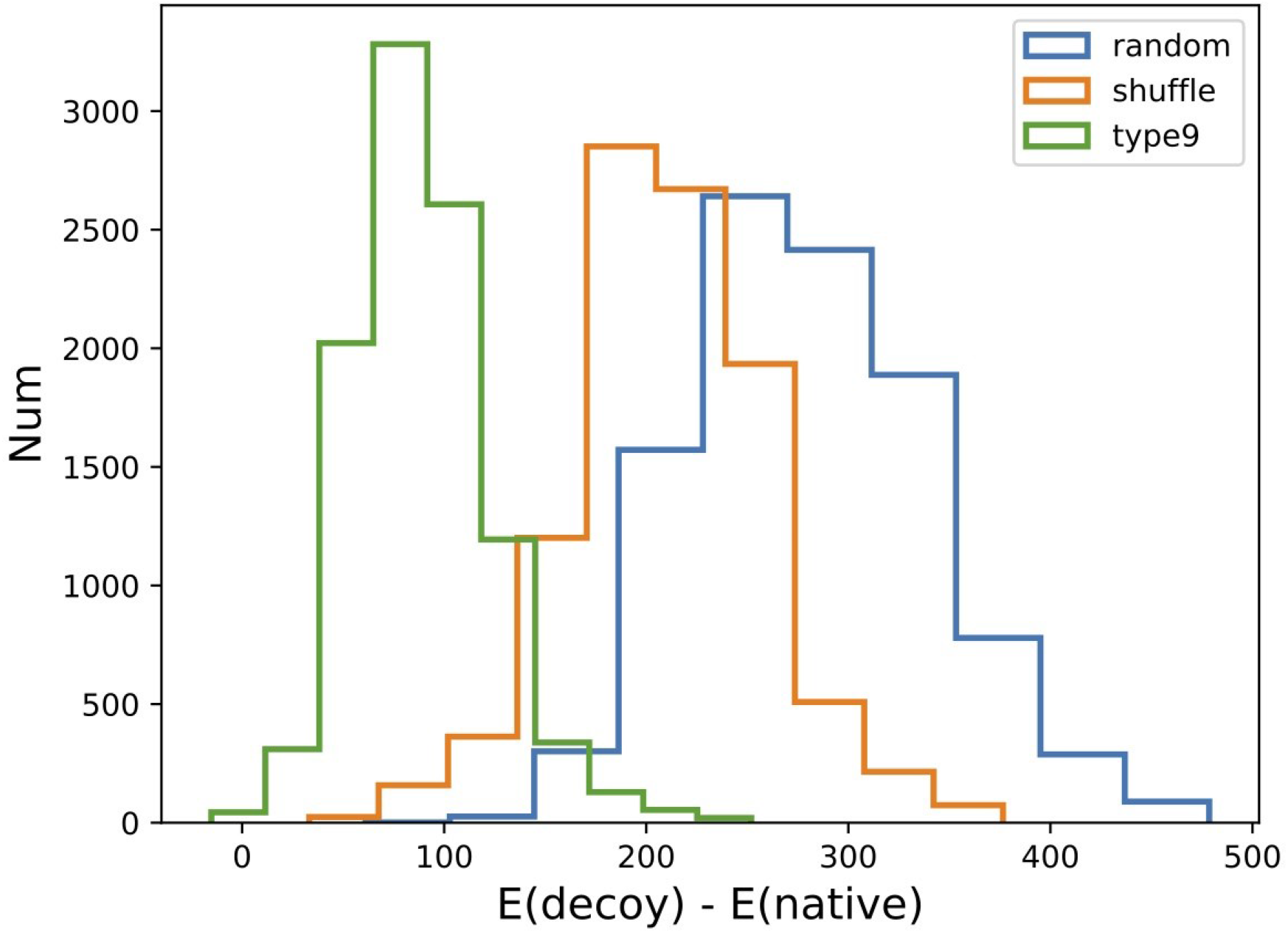
Distributions of energy differences E(decoy)-E(native) for three sequence decoy dataset. The decoy sequences were generated in three different ways: (1) substituting residues with chemically similar residues (green histogram), (2) shuffling the residues in the sequence (yellow histogram), and (3) mutating residues to random ones (blue histogram). The energies of the mutated sequences are higher than the native ones, and random mutations produce higher energies than mutations to similar amino acids

**Fig. S4.**
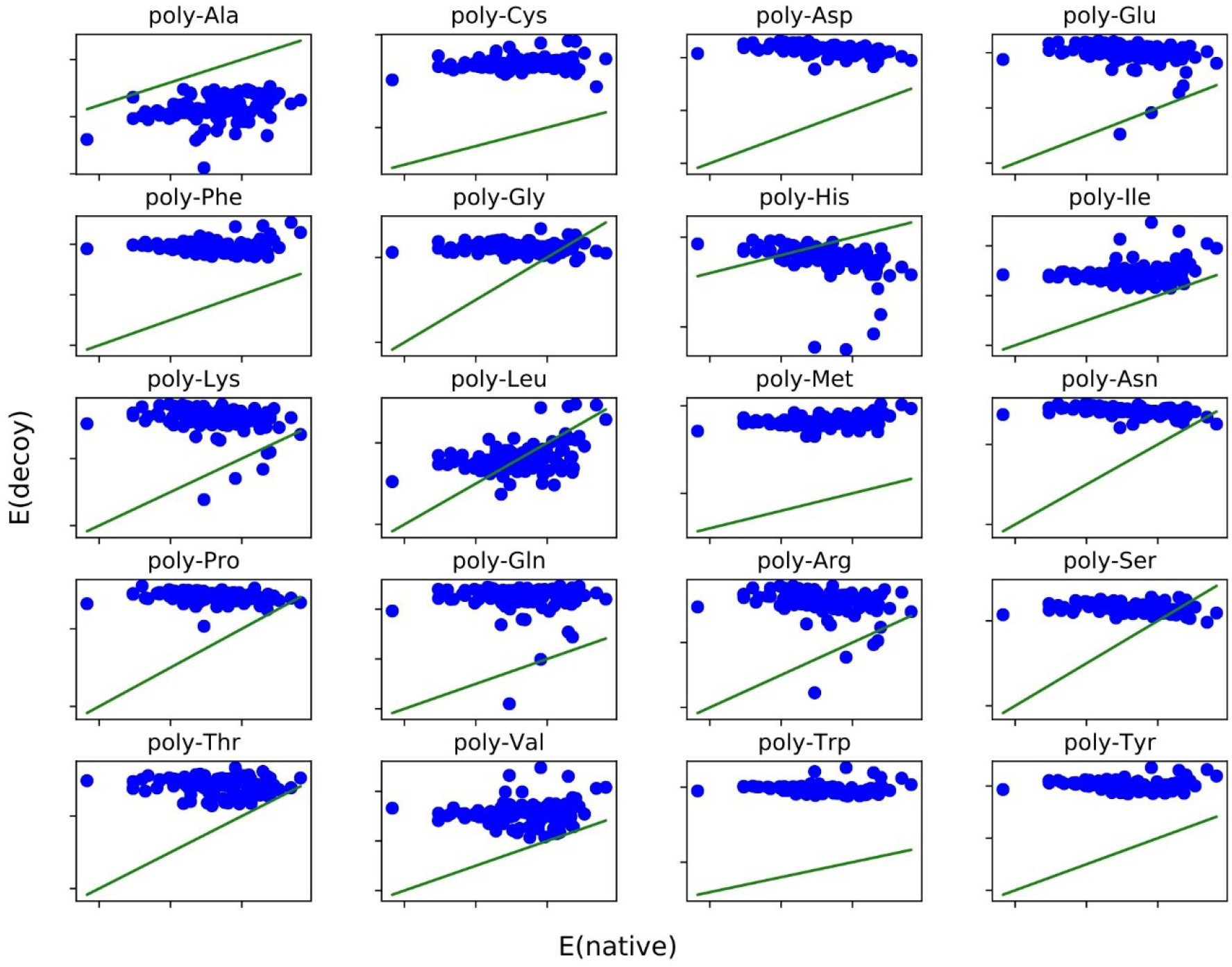
Comparison of the energies of native sequences and poly-X mutants (obtained by mutating every residue to the same one). In general, mutants have higher energies than native sequences, with the exception of poly-alanine decoys that systematically have lower energies compared to the respective native sequences. The same happens, to a lesser extent, also to poly-leucine and poly-histidine decoys.

**Fig. S5.**
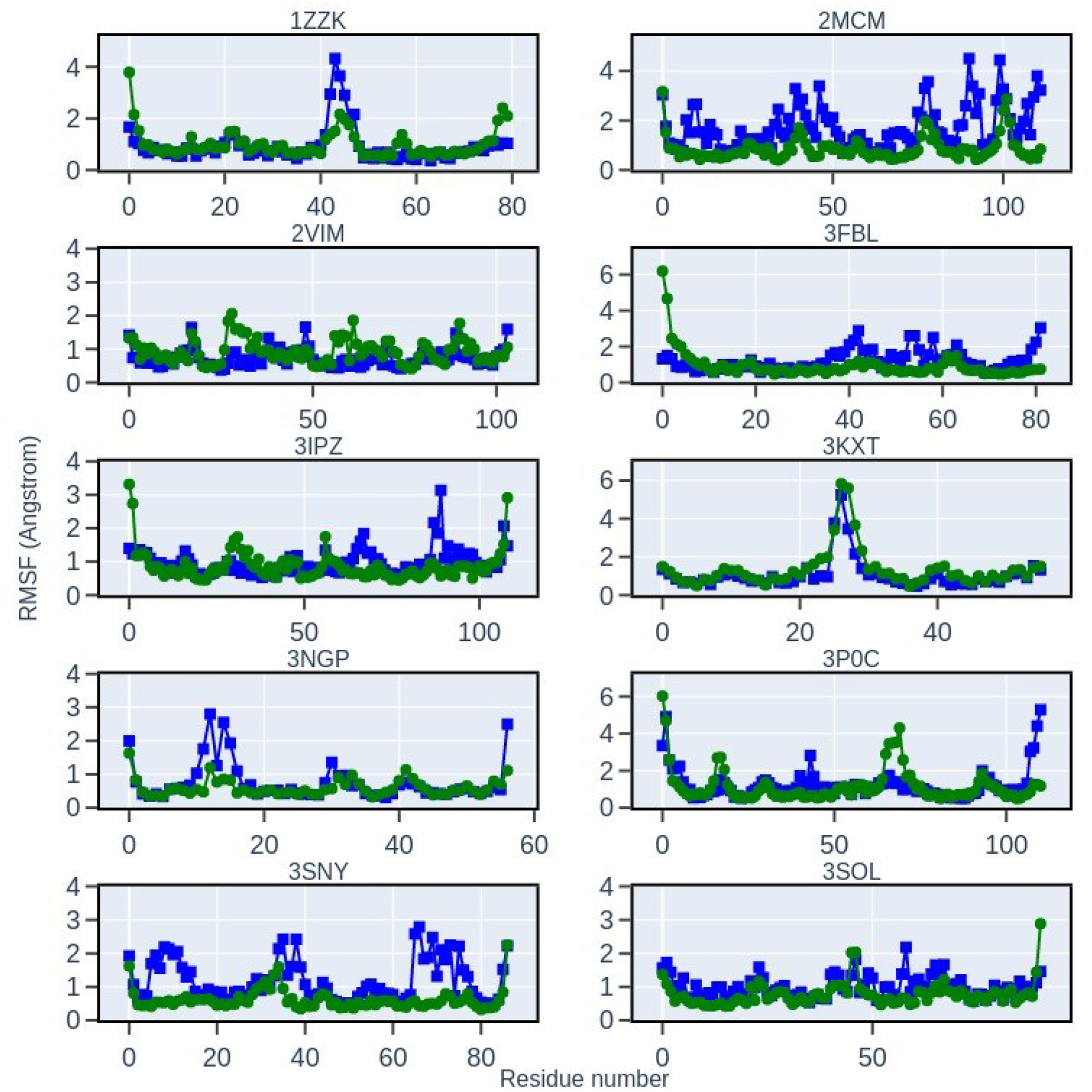
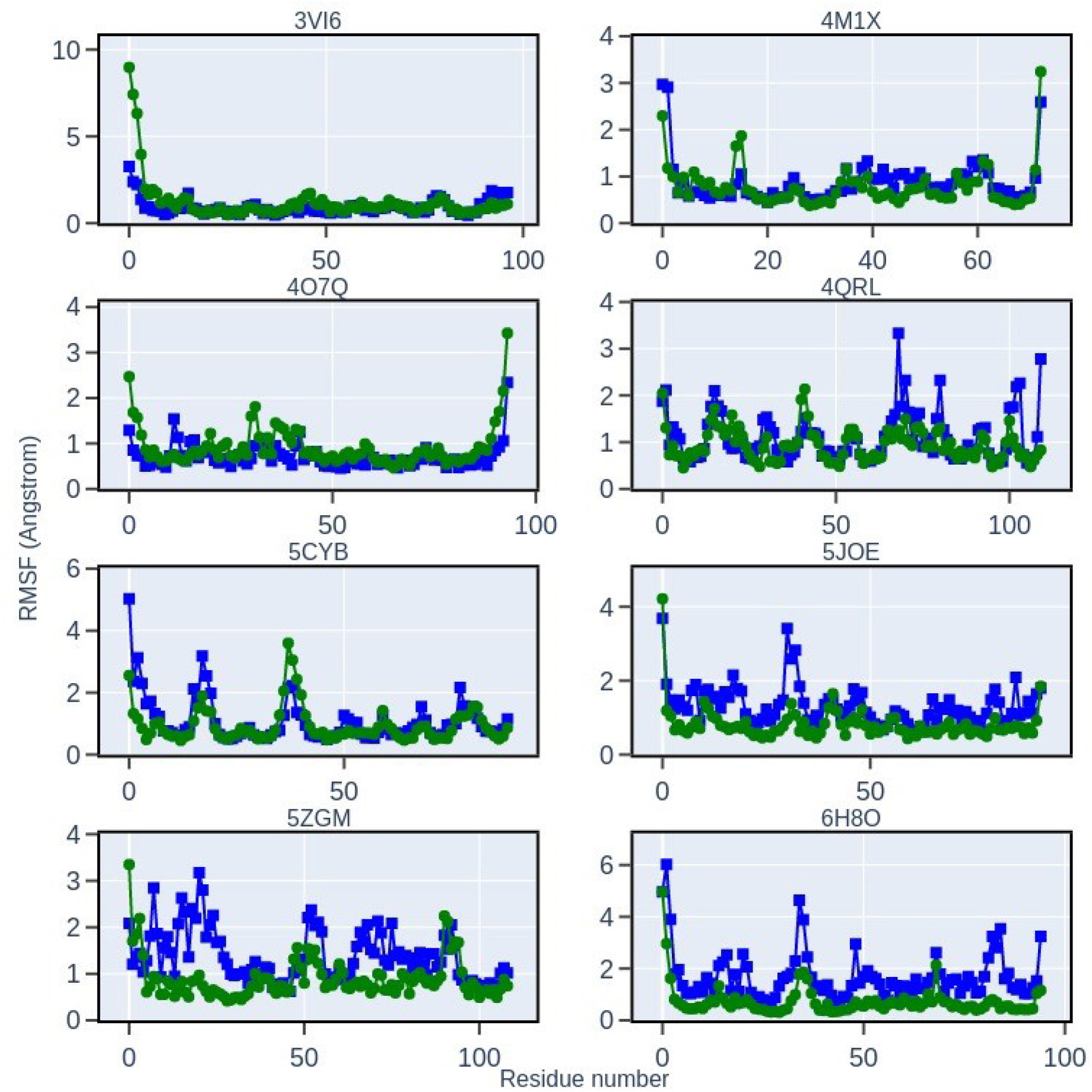
Comparison of the RMSFs for dynamic trajectories for all the 18 proteins in the test sample. One trajectory is generated by Langevin dynamics sampling of the NNEF (RMSF shown as blue squares). The other reference trajectory is generated from classical molecular dynamics simulations using *amber14SB* force field (RMSF shown as green dots).

**Fig. S6.**
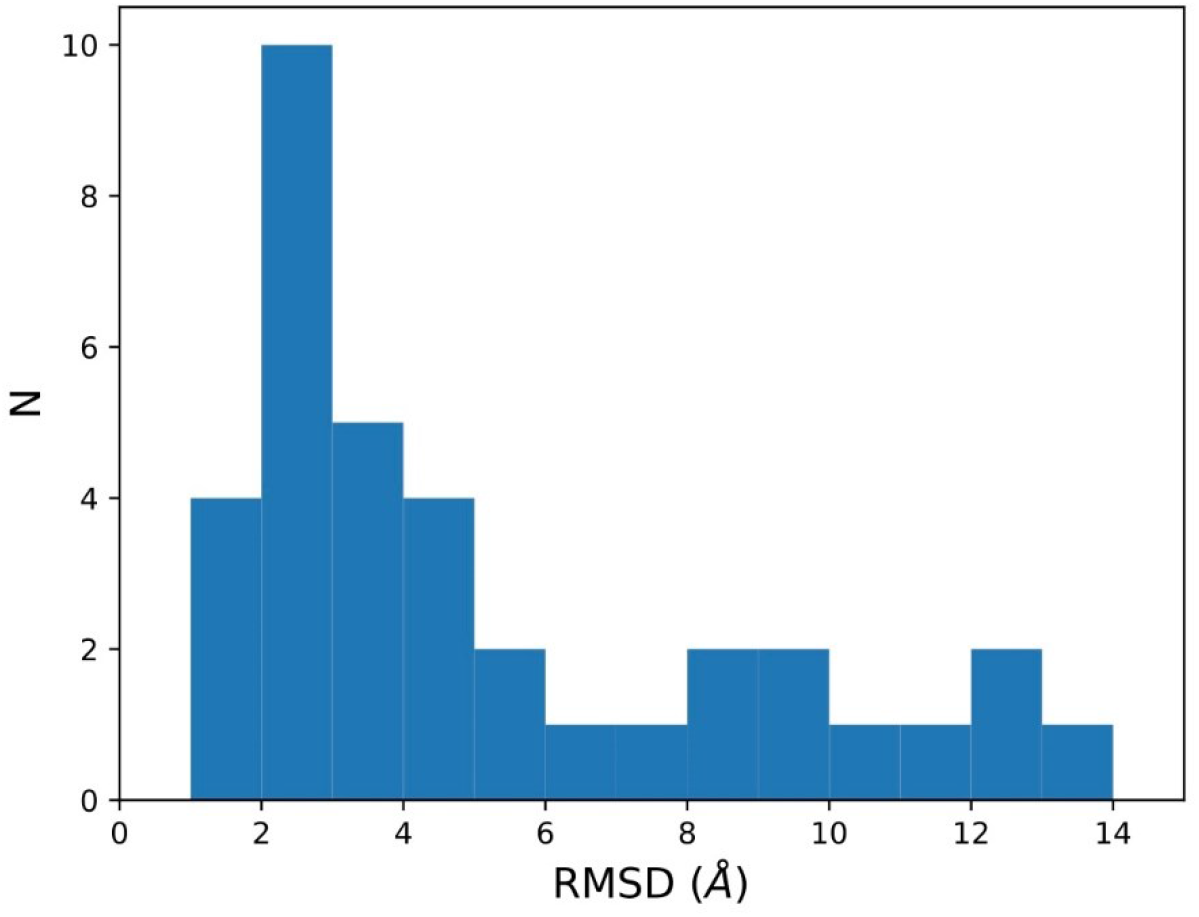
The distribution of RMSD between the predicted structure using *TrRosetta* and the target structure for 36 designed proteins. For about two thirds of the designed sequences, the predicted structures match the target structures with RMSD < 5.0Å.

## Notes

### Competing Interest Statement

The authors have declared no competing interest.

